# Quantifying transport in crowded biochemical environments

**DOI:** 10.1101/014704

**Authors:** Ruth E. Baker, Matthew J Simpson

## Abstract

Transport of cells and biochemical molecules often takes place in crowded, heterogeneous environments. As such, it is important we understand how to quantify crowded transport phenomena, and the possibilities of extracting transport coefficients from limited observations. We employ a volume-excluding random walk model on a square lattice where different fractions of lattice sites are filled with inert, immobile obstacles to investigate whether it is possible to estimate parameters associated with transport when crowding is present. By collecting and analysing data obtained on multiple spatial scales we demonstrate that commonly used models of motility within crowded environments can be used to reliably predict our random walk data. However, infeasibly large amounts of data are needed to estimate transport parameters, and quantitative estimates may differ depending on the spatial scale on which they are collected. We also demonstrate that in models of crowded environments there is a relatively large region of the parameter space within which it is difficult to distinguish between the “best fit” parameter values. This suggests commonly used descriptions of transport within crowded systems may not be appropriate, and that we should be careful in choosing models to represent the effects of crowding upon motility within biochemical systems.

---

Crowding can arise from geometrical restrictions and mutual obstruction, both of which are relevant to the transport of cells and biochemical molecules [13, 14]. For example the interior of the living cell is highly structured with both mobile and immobile molecules [15]. The presence of obstacles in the intracellular and intercellular environments leads to nonspecific interactions that can have a significant influence on the macroscopic transport properties in these systems. Specific examples of biological processes that have been shown to display sub-diffusive behaviours as a result of crowding include the motion of lipid granules in living yeast fission cells [10], protein diffusion in aqueous solution [26], motion of proteins and oligomers on membrances [7, 11], diffusion of Golgi membrane proteins in the endoplasmic reticulum and Golgi apparatus [36, 37], and receptor diffusion within the plasma membrane [25].

Given that these types of environments are ubiquitous throughout biology [3, 7], understanding how to quantify, parameterise and predict transport processes taking place in crowded environments is a major challenge relevant to our understanding of many biochemical systems. To this end, a number of computational studies have been undertaken to try and elucidate the effects of macromolecular crowding upon the myriad of reaction-transport processes that govern the behaviour of biochemical systems. Examples include the study of how crowding and confinement influences chemical and biochemical reactions in complex, crowded biological tissues [15, 16, 24, 34], crowded membrane protein dynamics [20, 25, 36], and the study of enzyme kinetics in crowded media [18, 22].

Many mathematical models of transport (and reaction) in crowded media have also been explored. In general, two main frameworks are used. For the study of single molecule motility, the mean squared displacement (MSD) is generally assumed to obey the generalised Einstein-Smoluchowski equation^1^

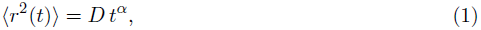

where the generalised diffusion coefficient, *D*, has units of L^2^ T^−*α*^, and *α* is a measure of the deviation from “normal” diffusion (*α* = 1). In general, the effects of crowding upon motility are assumed to give rise to values of *α* ϵ (0,1) [13, 14].

On the population-level, a generalised transport equation of the form

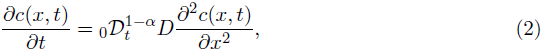

is often employed to describe how the density of the species under consideration evolves with time [1, 32, 13]. Here, the Riemann-Liouville operator, 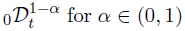, is defined through the relation

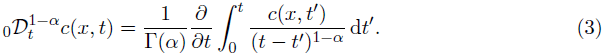

Equation (2) can be derived using a continuous-time random walk approach that describes the probability of finding a particle at position *x* at time *t*, given a probability density function that defines the jump length and time between successive jumps [13, 14]. Such frameworks have been used to explore subdiffusion-limited reactions [39, 40], morphogen gradient formation [41], the geminate recombination problem [30], chemotaxis and propagating fronts [4, 5] and Turing pattern formation [35].

However, many modelling studies employ equations (1) and (2) without careful consideration of whether they are appropriate for the system of interest. Further, the question of how much data is sufficient to characterise the nature of transport processes taking place within a crowded environment has not been, to our knowledge, carefully documented within the literature. Our study attempts to fill this gap. We explore two questions: firstly are equations (1) and (2) suitable individual- and population-level descriptions of transport within crowded media; and secondly, how much data is required to accurately estimate the parameters *D* and *α* that quantify transport processes taking place in crowded media.

To this end, we explore transport within a crowded environment where the crowding is described by the simplest possible mechanism; a population of randomly located immobile obstacles on a lattice [8, 9, 27, 29, 33]. Employing stochastic simulation techniques, we characterise the motion of a single random walker (known hereafter as an “agent”), which could represent the motion of a cell or molecule, as it moves through different crowded environments over different timescales. We use estimates of the MSD to characterise the motion using the diffusive framework of equation (1). Results of this type were first presented by Saxton [27, 28]: he demonstrated how the MSD of a single random mobile agent (known as a “tracer” agent) deviated from the usual linear relationship with time, *⟨r^2^(t)⟩* = *Dt^*α*^* with *α* = 1, as the extent of crowding increased. In addition to characterising the MSD, we use results from repeated simulation of our model to construct an estimate of the spatial distribution of tracer positions at various times. This distribution is characterised using the general diffusive framework of equation (2) and results compared to the information on motion gained from estimates of the MSD.

To provide a complementary description of the motion of a population of such agents, we also consider simulations of the collective motion of a population of agents as the population moves through similarly crowded environments. At most one mobile agent or immobile obstacle is allowed per site, in the same manner as for exclusion processes [31]. In this case crowding effects arise from both the immobile obstacles and the population of mobile agents themselves (self-crowding). Using stochastic simulation algorithms we construct ensemble-averaged agent density profiles for the collective motion case. We illustrate how the formation of a concentration gradient may be affected by obstacles in the microenvironment. This kind of data, and our ability to parameterise it using the general diffusive framework of equation (2), allows us to examine the relationship between microscopic, single-agent descriptions of the motion with a macroscopic description of the motion of a population of such agents moving through equivalently crowded environments.

In addition to examining the relationship between the description of single agent transport and collective transport in crowded environments, we also examine how our ability to characterise motion through crowded environments is affected by the availability of experimental data. We achieve this by studying how our parameter estimates are affected by having access to different numbers of realisations of the same stochastic process.

In summary, we show that quantitative descriptions of transport through crowded environments differ depending on whether we examine the process from the point of view of a single tracer agent or whether we consider the collective motion of a population of agents through a crowded environment. Furthermore, our results indicate that, although equations (1) and (2) can be used to reliably predict data on motility within crowded environments, extremely high quality data is required to identify the parameters *D* and **α** describing the process. These observations are significant since many experimental descriptions of crowded biochemical transport rely on having access to a relatively small number of experimental trajectories to estimate the MSD of tracer agents. Further when crowding is present, even with unfeasibly large amounts of data, there is a relatively large region of parameter space within which it is difficult to distinguish between the best-fit parameter values. Given that this problem does not arise in the non-crowded case, we suggest that commonly used descriptions of motility within crowded systems may be inappropriate. This means that we should be careful in choosing models to represents the effects of crowding in biochemical systems.

The outline of this work is as follows: in Methods we outline the algorithms used to generate our discrete data and give a quantitative characterisation of the different systems we consider; in Results we discuss results concerning both single-agent and population-level data; and in Discussion we briefly discuss our results and their implications for modelling transport in crowded biochemical systems.

## Results

The main aim of our study has been to investigate a range of simple volume-excluding systems, and investigate how individual- and population-level behaviours are affected by the presence of non-motile obstacles in the local environment. Our focus is on understanding the extent to which we can quantify motility in crowded environments, given different types and amounts of data.

### Quantifying motility using mean squared displacement data

We first investigated how the ensemble-averaged MSD of a tracer agent increases with time as the background obstacle concentration changes. Our aim was to understand: (i) how increases in obstacle concentrations affect the MSD; and (ii) to what extent differing amounts of data can be used to reliably characterise motility under crowded conditions.

#### Increases in obstacle density result in decreases in mean squared displacement

As expected, increases in the obstacle density result in decreases in the ensemble-averaged MSD. In particular, in Figure 2 we show how the ensemble-averaged MSD of a tracer agent increases with time, for a range of obstacle concentrations increasing from 0% to 50% in increments of 5%. The top plot shows the relationship between MSD and time, indicating, as expected, that the MSD increases less rapidly with time as the obstacle density is increased. These results are consistent with other work, such as that presented by [27, 28]. The lower plot shows log_10_(⟨*r*^2^⟩/*t*) as a function of log_10_(*t*). In this case, assuming ⟨*r*^2^(*t*)⟩ = *Dt*^*α*^, we could potentially use this plot to give estimates of *D* and *α* by considering the intercept (log_10_(*D*)) and the gradient (*α* — 1). Here normal diffusion (*α* = 1) is indicated by a straight line with zero gradient. Once again, as suggested by [27], we see a range of behaviors with, for several blockage densities, a transition between anomalous (*α* < 1) and normal (*α* = 1) diffusion as time increases. In fact, for all the obstacle densities shown in Figure 2, the curves will eventually show a transition between anomalous and normal diffusion as all obstacle densities are below the percolation threshold for a 2D square lattice (approximately 59.27% [19]). However, over the time scale illustrated in Figure 2, the systems with obstacle densities of 40%, 45% and 50% have not yet reached the normal diffusion regime.

**Figure 1:**
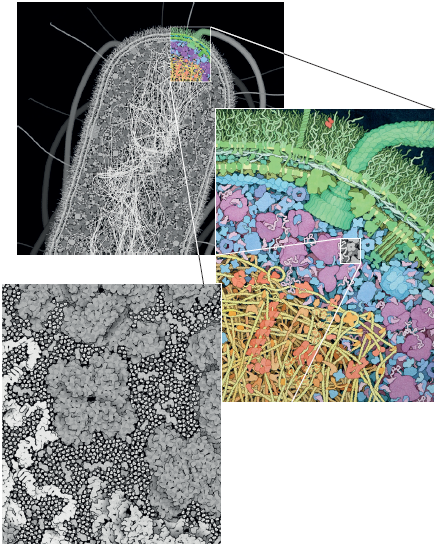
An illustration of the crowded nature of biological systems. The sequence of magnified images shows a portion of an *Escherichia coli* bacterium, including all macromolecules. The colour scheme differentiates between different functional compartments of the cell, including the cell wall and flagellar motor (green), soluble enzymes and other proteins (turquoise), ribosomes and tRNA (magenta), DNA (yellow) and proteins in the nucleoid (orange). Reproduced with modification from Machinery of Life, Copernicus Books, Springer, Figures 4.1–4.3, D. S. Goodsell, 2009 [permission pending].

**Figure 2.**
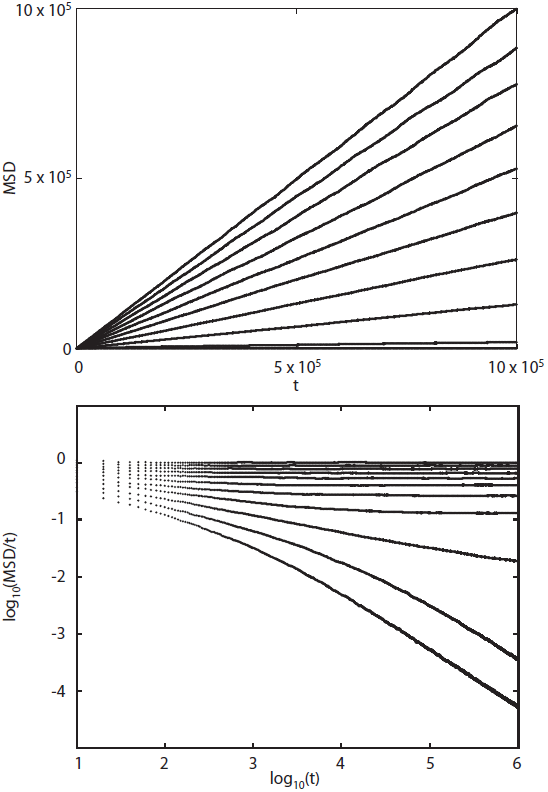
Variation in the MSD of a tracer agent with varying obstacle concentrations, ranging from 0 — 50% in increments of 5%. The MSD decreases monotonically as a function of obstacle concentration. Top: a plot of MSD, (*r*^2^), as a function of time, *t*. Bottom: a plot of log_10_〈*r*^2^〉/*t*) against log_10_(*t*). The discrete simulations were carried out as described in detailed in Methods on a 1000 × 1000 square lattice and results are averaged over 10^4^ realisations.

#### Infeasibly large amounts of data are required to quantify the impact of crowding upon diffusion

We attempted to estimate *D* and **α** directly from data on the MSD as a function of time (Figure 3). Note that we constrained our estimation algorithms to consider 0 < *D* ≤ 0.25 and 0.5 ≤ **α* ≤* 1. This range of *D* is relevant since *D* = 0.25 is an upper bound for our random walk model when there are no obstacles present [13]. While, in principle, we could have restricted our parameter estimation algorithms to 0 < *α* ≤ 1, preliminary results (not shown) indicated that the best fit estimates always corresponded to the smaller region, 0.5 ≤ **α* ≤* 1. Furthermore, our results suggest that focusing on this smaller region, 0.5 ≤ **α* ≤* 1, always contained the best fit parameter values for all problems considered. Going from the top to bottom rows in Figure 3 we present results from simulations with obstacle densities of 0%, 20%, and 40%. In the left-most column we show ensemble-averaged MSD results generated using different numbers of realisations. Error bars indicate one standard derivation from the mean. We see that for *N* = 10 and *N* = 100 realisations the error bars are large and there are significant differences between the averaged MSD curves for the same obstacle density. These plots indicate that very large amounts of data (*N* >> 100) are required to provide reliable estimates of the motility parameters (*D*, *α*) from ensemble-averaged MSD data over intermediate time scales, a result in line with that of an experimental study carried out by Qian et al. [23].

**Figure 3:**
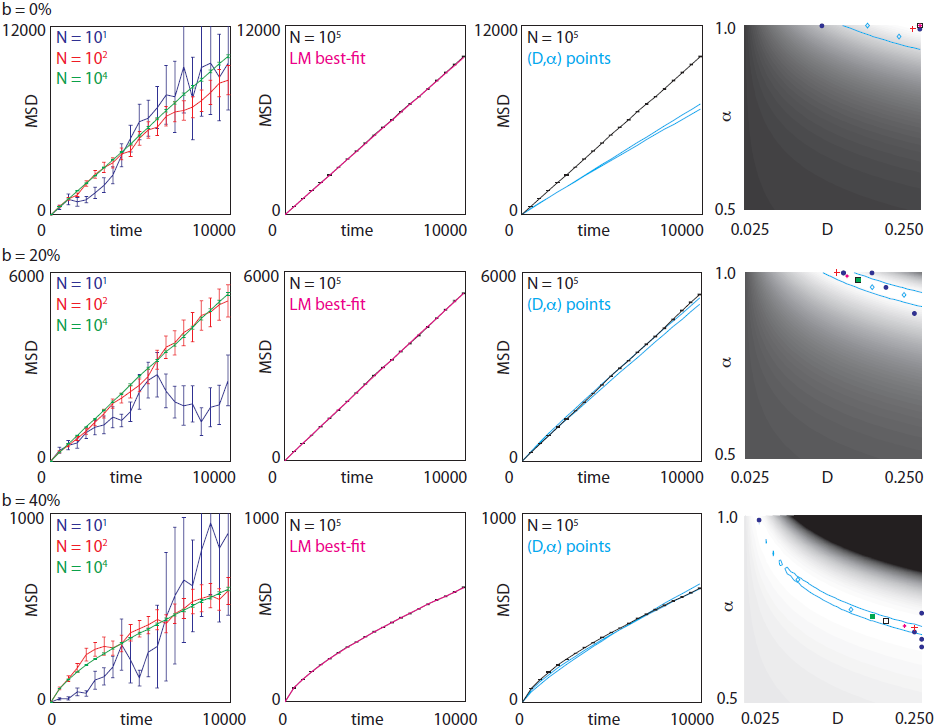
Ensemble-averaged MSD data with different obstacle densities (*b*). Column 1: Ensemble-averaged MSD curves obtained using different numbers of realisations, *N*. Error bars indicate one standard deviation from the mean. Column 2: Ensemble-averaged MSD using *N* = 10^5^ realisations, compared with plots of ⟨*r*^2^⟩ = *Dt^α^* where values for *D* and *α* were estimated using the Levenberg-Marquardt fitting algorithm. Column 3: Ensemble-averaged MSD from *N* = 10^5^ realisations, compared with plots of ⟨*r*^2^⟩ = *Dt^*α*^* where the *(D*, *α*) parameter combinations are indicated by the light blue diamonds on the corresponding plot in Column 4. Column 4: Sum-squared difference between ensemble-averaged MSD data using *N* = 10^5^ realisations at *t* = 10,000 and the curve ⟨*r*^2^⟩ = *Dt^*α*^* for various values of *D* and *α*. The greyscale indicates the sum-squared difference, *ϕ*(*D*, **α*; t*) as defined in equation (5), for each (*D*, *α*) pair with white being low and black being high. The blue lines enclose a region containing the fifty points with the smallest *ϕ*(*D*, *α*; *t*). Dark blue dots - minimum ϕ(*D*, **α**; *t*) from five different ensemble-averaged MSD data sets each generated with *N* = 10; red cross - minimum *ϕ*(*D*, **α**; *t*) from data averaged over *N* = 10^2^ realisations; green square - minimum *ϕ*(*D*, **α**; *t*) from data averaged over *N* = 10^4^ realisations; black square - minimum *ϕ*(*D*, *a*; *t*) from data averaged over *N* = 10^5^ realisations; pink star - Levenberg-Marquardt estimate; light blue diamonds - two randomly chosen points within the “minimum region”. The discrete simulations were carried out as described in detailed in Methods.

However, we were able to find values of (*D*, *α*) that gave good agreement between the predicted power law MSD of equation (1) and ensemble-averaged MSD data when the number of realisations was high. In the second column from the left in Figure 3 we show the results of using the Levenberg-Marquardt algorithm to fit equation (1) to ensemble-averaged MSD data generated using *N* = 10^5^ realisations. In each case, the MSD curve, equation (1), plotted using the values of *(D,*α*)* predicted using the Levenberg-Marquardt algorithm is very close to the ensemble-averaged MSD curve.

For data of decreased quality the predictions for (*D*, *α*) are highly variable. The rightmost plots in Figure 3 indicate parameter estimates from a range of other data. The grey shading shows the sum-squared difference, *ϕ(D, *α*; t)* (see equation (5) of Methods), between MSD ensemble-averaged data using *N* = 10^5^ realisations and the curve ⟨r^2^⟩ = *Dt^*α*^* for various values of *D* and **α**. The region enclosed by the light-blue lines indicates the fifty *(D,*α*)* pairs with the smallest sum-squared difference, ϕ(*D, *α*; t*), and the black, hollow square indicates where the minimum value of *ϕ(D, *α*; t*) is attained (note that the error surface was generated by considering 11 equally-spaced values of *D* and 51 equally-spaced values of *α*). The small pink dot indicates the best-fit parameter combination as estimated using the Levenberg-Marquardt algorithm. Note that it is not always coincident with the minimum of the sum-squared difference, *ϕ(D, *α*; t*). This is because we have only evaluated *ϕ(D, *α*; t*) using a discrete number of *(D, *α**) pairs and also potentially due to the Levenberg-Marquardt algorithm finding a local minimum of *ϕ(D, *α*; t*) (results are sensitive to the initialised values of *(D, *α*)* and also the algorithm parameters). The red cross and filled green square indicate the minimum sum-squared difference for data averaged over 10^2^ and 10^4^ realisations, respectively, and the five large blue dots indicate the minimum sum-squared difference obtained from five different ensemble-averages using *N* = 10 realisations. We chose to present data for *N* = 10 realisations since this corresponds to a realistic amount of data that could be achieved in a real biological experiment. For each of the three rightmost plots we picked two random points from inside the “minimum region” (containing the 50 (*D, *α**) pairs with the smallest *ϕ(D, *α*; t))* and they are indicated by light-blue diamonds. In the third column from the left we plot ⟨r^2^(t)⟩ = *Dt^*α*^* for these values of *(D, *α*)* and compare the plots to ensemble-averaged MSD data from 10^5^ realisations. This gives us some insight into the sensitivity of the predictions of equation (1) to the parameters.

For the case of no crowding (0% obstacles), ensemble-averaged MSD data generated using at least *N* = 10^2^ realisations gives estimates of *(D, *α*)* close to those predicted theoretically: *(D, *α*)* = (0.25,1.00) [13, 21]. However, with *N* = 10 realisations results vary significantly. In particular, upon increasing the range of values of *α* to *α* ∈ [0.5,1.2] our Levenberg-Marquardt algorithm estimates *(D, *α*)* pairs with minimum sum-squared differences that differ wildly from *(D,*α*)* = (0.25,1.00) (results not shown). However, we note that the *ϕ(D, *α*; t)* surface (plotted in Figure 3 and generated using *N* = 10^5^ realisations) has a well-defined minimum: plotting *(r^2^(t))* = *Dt*^*α*^ for two randomly chosen points within the minimum region gives results that deviate significantly from the *N* = 10^5^ data. This indicates that it is relatively straightforward to accurately determine the transport parameters when no obstacles are present *(b = 0%)*, given a sufficient amount of data.

When obstacles are present the results are more difficult to interpret. As the obstacle density increases, the minimum region moves away from the top-right-hand corner of the sum-squared difference plot and occupies a curved region that encompasses a range of values of *(D, *α*)*. The *(D, *α*)* pair with the minimum sum-squared difference is often significantly different from the *(D, *α*)* pair predicted by the Levenberg-Marquardt algorithm, although both tend to lie in the minimum region. The deviations in predictions for *(D, *α*)* are more pronounced for data averaged over fewer numbers of realisations, as expected. Finally, the sum-squared difference surface has a less well-defined minimum: in each case the randomly chosen *(D,α)* pairs in this region give rise to MSD curves given by equation (1) that are close to those predicted from the *N* = 10^5^ data. This indicates that there are many possible choices of *(D, *α*)* that match the observed data reasonably well when *b* > 0, and suggests that the power law of equation (1) may not be an appropriate description of motility in crowded situations. Note that this is in contrast to the obstacle-free case in which only values of *(D, *α*)* very close to the well-defined minimum provide a good fit to the data.

In summary, our individual-level results show that estimates of *(D, *α*)* are very sensitive to the amount of trajectory data that is available. In particular, a very large number of trajectories need to be considered before any kind of reliable predictions can be made from data. These results are consistent across data fitted over different time intervals (note that in the Supplementary Material we show additional results with the same analyses conducted at times *t* = 1,000 and *t* = 5,000, and the results and conclusions are unchanged). Moreover, we observe that only for unobstructed diffusion (no obstacles, b = 0) is there a well-defined minimum to the sum-squared difference surface. This implies that for obstructed diffusion there is always a range of *(D, *α*)* pairs that fit the data well. These results have significant implications in terms of our ability to characterise motility in crowded environments, and the validity of equation (1) to describe MSD data therein.

#### Quantifying motility using single-agent density data

We next considered how our ability to characterise motility changes when we instead consider single-agent averaged density data. We used multiple realisations of a model system in which a single tracer agent undergoes a random walk on a crowded lattice to estimate the agent density as a function of space and time. We investigated how the average density profile varied as a function of obstacle density, and then tried to quantify this variation by fitting the density profile to the solution of a generalised transport model, given by equation (2). Our results are summarised in Figure 4 where we show plots similar to those in Figure 3 but now with population-level agent density data. As in Figure 3, in subsequent rows of Figure 4 we present data collected from realisations with obstacle densities of 0%, 20%, and 40%, respectively.

**Figure 4:**
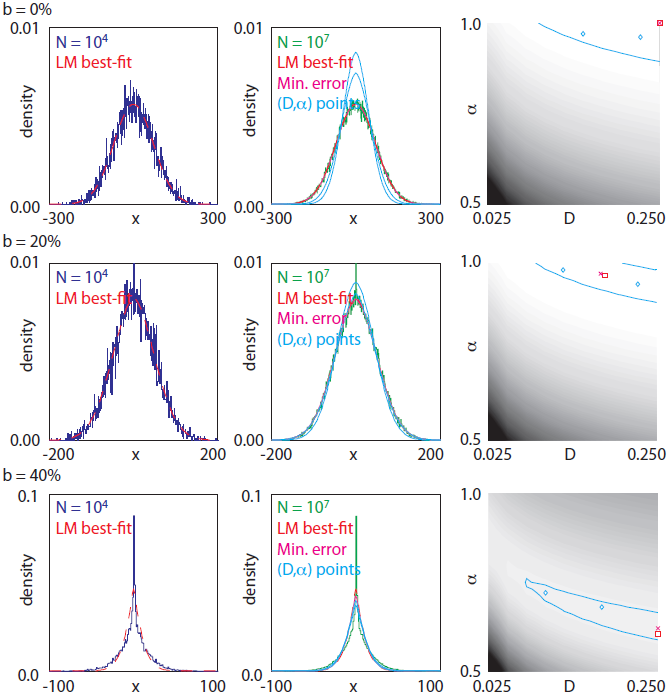
Ensemble-averaged density data with a range of obstacles densities (*b*). In each realisation, a single tracer is placed at a lattice site with *x* = 0 and a random y position and its position recorded at time *t* = 10,000. Averaging over a number of realisations gives a density profile. Column 1: comparison of results obtained from averaging over 10^4^ realisations and the best-fit solution of equation (2) obtained using the Levenberg-Marquardt algorithm fitted to data averaged over 10^7^ realisations. Column 2: results from averaging over 10^7^ realisations and density profiles generated using equation (2) with the Levenberg-Marquardt-estimated *(D,a),* the *(D,a)* parameter combinations indicated using blue diamonds in Column 3, and the (*D*, **α*)* pair at the minimum of the sum-squared difference plot in Column 3. Column 3: sum-squared difference between density data averaged over 10^7^ realisations at *t* = 10, 000 and the PDE solution for various values of (*D, *α**). The greyscale indicates the sum-squared difference with white being low and black being high. The blue region encloses the fifty closest points to the minimum. Red square - best-fit parameter values obtained using the Levenberg-Marquardt algorithm fitted to the data averaged over 10^7^ realisations; pink cross - minimum of the sum-squared error; blue diamonds - two randomly chosen points within the minimum region. The discrete simulations were carried out as described in detailed in Methods.

Averaging over *N* = 10^4^ realisations of the discrete processes gave rise to relatively noisy density profiles, though it was possible, for each obstacle density, to find parameters *(D, *α*)* for which the generalised transport model, equation (2), compared well with the discrete data. The left-most column shows density data collected from averaging over *N* = 10^4^ realisations of the discrete process compared with a density profile whose evolution is described by the generalised transport model, equation (2). The parameters *(D, *α*)* of the generalised transport model were estimated using the Levenberg-Marquardt algorithm with averaged discrete data obtained from *N* = 10^7^ realisations. In the right-most column we show the sum-squared difference between the density profile estimated from numerical solution of equation (2) (using the same range of *(D, *α*)* pairs as in Figure 3) and ensemble-averaged discrete data. The grey shading indicates the sum-squared difference between the density profile averaged over *N* = 10^7^ realisations and numerical solution of equation (2). The pink cross indicates the minimum sum-squared difference over this surface whilst the red square indicates the (*D*, *α*) pair estimated using the Levenberg-Marquardt algorithm. In all cases there is good agreement between the two estimates, however, the quality of the agreement is reduced when the ensemble data is averaged over fewer realisations (data not shown). The blue line demarcates minimum region (enclosing the fifty *(D, *α*)* pairs with sum-squared differences closest to the minimum sum-squared difference), and the blue diamonds indicate two randomly chosen points within this region. The data shows a similar qualitative trend to the individual-level MSD data with the minimum region encompassing a range of (*D, *α**) combinations and lying in a similar region of parameter space.

We then investigated the sensitivity of our predictions of the density profile to the values of (*D, *α*)*. We took two randomly chosen (*D, *α**) pairs within the minimum region and plotted the density profiles predicted by equation (2) using these parameter pairs (middle column of Figure 4). The same trend as for the individual-level data is seen in terms of the error surface having a well-defined minimum: for 0% obstacles the two randomly chosen points within the minimum region give rise to density profiles that deviate significantly from the averaged discrete data indicating that it is relatively straightforward to recover the true transport parameters. Alternatively, the density profiles for the 20% and 40% obstacle cases are far less sensitive to the choice of *(D, *α*)* and we observe that different combinations of *(D, *α*)* produce reasonable matches to the observed data.

Once again, our data illustrate that one must be very careful in trying to estimate the transport coefficients D and *α* from data. Relatively large quantities of data are required before one can make sensible estimates, with many orders of magnitude worth of data than is usually available from experiments necessary. Moreover, whilst the estimates obtained from MSD and density data show the same qualitative trends as the number of obstacles is increased, there are significant quantitative differences in the estimates even with high-quality data^2^.

#### Quantifying motility using multi-agent population-level data

Finally, we explored systems in which there is more than one tracer agent present, in order to understand the additional impact of self-crowding upon motility. These kinds of multi-agent simulations are relevant to many biological experiments which focus on collective migration where self-crowding are important [38]. To explore mult-agent population-level data we performed a range of simulations in which columns of the lattice with 1 ≤ *x* ≤ 20 were permanently filled with agents. If an agent leaves a site in this region at any time step of the algorithm, we immediately re-fill the vacant site. This means that at all times, for 1 ≤ *x* ≤ 20, *b*% of sites are filled with obstacles, and the remaining (100 — *b*)% of sites are filled with mobile agents. Given the remainder of the lattice is initially empty (except for obstacles), we see a gradual gradient in agent density build up across the lattice over time (see Figure 5), as expected.

**Figure 5:**
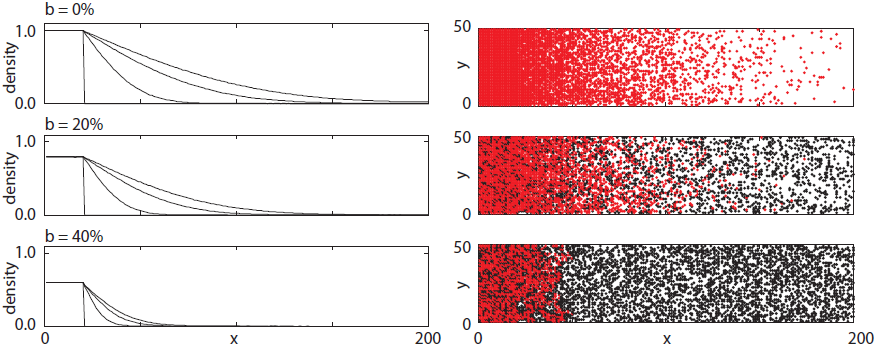
The effect of crowding on the formation of concentration gradients for different obstacle percentages (*b*). In each case, the left-hand plots show the results of averaging over 1000 realisations of the discrete process at *t* = 0, 1,000, 5,000, 10, 000, whilst the right-hand plots show the results of one realisation at *t* = 10,000. The red (grey) dots indicate motile agents whilst the black dots indicate obstacles. The discrete simulations were carried out as described in detailed in Methods.

To quantify the impact of both obstacles and self-crowding upon motility, we carried out similar analyses as for the single-agent density study. Our results indicate that for 0% obstacles with sufficient data it is possible to accurately predict theoretically estimated values of *(D, *α*)* = (0.25,1.00) and that the sum-squared difference surface has a well-defined minimum (Figure 6, top row). Sensitivity to the choice of (*D*, *α*) is clear since the solutions of equation (2) using the two randomly chosen (*D, *α**) pairs within the minimum region are very different to the observed data. This result is consistent with the individual-level and constant density population-level results presented previously. Again, when obstacles are present in the system, the sum-squared difference surface has a less well-defined minimum, and a range of (*D, *α**) parameter combinations are able to accurately predict population-level data (Figure 6, middle and bottom rows). This insensitivity to the choice of (*D, *α**) is demonstrated by the fact that solutions of equation (2) using two randomly chosen points within the minimum region are practically indistinguishable from the observed data, showing that many possible choices of (*D, *α**) give comparable solutions.

**Figure 6:**
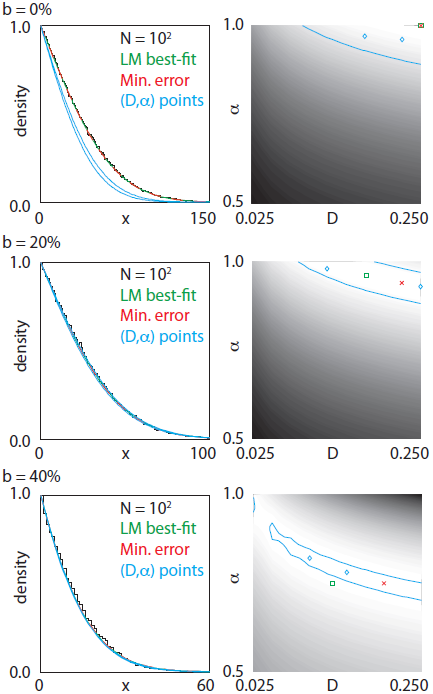
Gradient density data collected from multiple realisations with 0% (top row), 20% (middle row) and 40% (bottom row) obstacles. Simulations were carried out in the same manner as in Figure 5 and then the density rescaled to be unity at the left-hand edge of the domain. Column 1: comparison of results obtained from averaging over 10^2^ realisations, the best-fit solution of equation (2) obtained using the Levenberg-Marquardt algorithm fitted to the data, the minimum of the error surface in Column 2 and two *(D, *α*)* parameter combinations indicated using blue diamonds in Column 2. Column 2: sum-squared error between density data averaged over 10^2^ realisations at *t* = 10, 000 and the PDE solution for various values of (*D*, *α*). The greyscale indicates the error with white being low and black being high. The blue region encloses the fifty closest points to the minimum. Green square - best-fit parameter values obtained using the Levenberg-Marquardt algorithm fitted to the data averaged over 10^2^ realisations; red cross - minimum of the sum-squared error; blue diamonds - two randomly chosen points within the minimum region. The discrete simulations were carried out as described in detailed in Methods.

In Figure 6 we show results obtained from using the Levenberg-Marquardt algorithm to estimate the parameters (*D, *α**) that give a best fit between our discrete data and the numerical solution of equation (2). In the left-hand column we show results from *N* = 10^2^realisations of the discrete algorithm. We observe that the spatial extent of the gradient decreases as the obstacle density increases. We also show results from solution of equation (2) with values of (*D, *α**) estimated using both the Levenberg-Marquardt algorithm and finding the minimum sum-squared difference between the ensemble-averaged data and numerical solution of equation (2). In the right-hand column of Figure 6 error plots similar to those shown in Figure 4 are given. The grey shading indicates the sum-squared difference between numerical solution of equation (2) and data averaged over *N* = 10^2^ realisations with the red cross indicating the minimum. The hollow green square indicates the parameter combination estimated using the Levenberg-Marquardt algorithm. Again, the blue lines demarcate the minimum region, the fifty (*D*, *α*) pairs with sum-squared differences closest to the minimum, and the blue diamonds two randomly chosen points in this region. Plots of the density profiles obtained using these values of (*D*, *α*) are also given in the left-hand column. Additional data evaluated for final times *t* = 1, 000 and *t* = 5,000 is shown in the Supplementary Material and gives rise to similar results.

In summary, our results confirm the conclusions made using MSD and single-agent density data: a very large amount of data must be collected before being able to make any kind of reliable parameter estimation and, further, that when obstacles are present in the system a range of (*D*, *α*) parameter combinations give a good fit to ensemble-averaged data.

## Discussion

The proper function of biochemical processes is controlled to a large extent by the transport of cells and biochemical molecules and hence, in general, it must be tightly regulated. In contrast to highly idealised *in vitro* experiments, these transport processes often take place in extremely crowded, heterogeneous environments where transport can be significantly affected by the specific nature of the local microenvironment [3, 15, 16]. The aim of this work has been to invoke a simplified mathematical description of transport within a crowded environment, with a view to understanding the extent to which it is possible to characterise transport in crowded systems. As such, we considered random walk data collected on a range of spatial scales, investigating how our ability to estimate transport coefficients of the system depend on the amount of data available and the spatial scale on which it is collected. We assumed, as our theoretical models, two commonly used (but often unvalidated) models: first we supposed that the MSD can be represented by a power law in the form of equation (1); and second that the population density can be modelled using a generalised transport equation such as (2).

We first presented MSD data collected from tracking the position of a single tracer agent as it underwent a random walk on a lattice with various proportions of lattice sites occupied by immobile obstacles. Our data demonstrates that a significant number of trajectories (orders of magnitude more than generally collected from typical biological experiments) must be recorded to obtain ensemble-averaged MSD data from which parameter estimates can be reliably made. Our results also show that the presence of obstacles in a system reduces the extent to which it is possible to reliably estimate transport parameters as the sum-squared difference surface generated using the ensemble-averaged data and theoretical prediction exhibits a less well-defined minimum. Furthermore, we observe that the minimum becomes increasingly less well-defined as the density of obstacles in the system increases. This result suggests that equation (1) may not be suitable for the systems that we investigate here, and so care must be taken when employing equation (1) as a description of crowding in biological systems. In fact, within the framework of our investigation, although diffusion appears anomalous *(*α* <* 1) on intermediate time scales, it becomes normal *(*α* = 1*) on long time scales (see Figure 2 and [2, 27]) meaning that a single value of *α* is insufficient to describe the MSD data over time scales of several orders of magnitude.

These individual-level results were substantiated by population-level data where we analysed data obtained both from simulations where the tracer agent was obstructed by a constant density of immobile obstacles, and by a combination of immobile obstacles and other motile tracer agents. In particular, we again see a reduction in the extent to which it is possible to reliably estimate transport parameters by fitting solutions of equation (2) to data as the sum-squared difference surface generated using ensemble-averaged data and the theoretical prediction does not exhibit a well-defined minimum. In the same vein as for the MSD data, this suggests that care should be taken when using equation (2) to describe transport within crowded biochemical systems. It is noteworthy, also, that although the individual- and population-level constant density experiments showed the same qualitative trends, parameter estimates obtained by fitting to a theoretical MSD curve and a generalised transport equation did not give the same quantitative estimates.

In summary, the take-home message from our study is two-fold. First, on intermediate time scales, equations (1) and (2) can provide a good fit to individual- and population-level data on motility in crowded systems. However, when obstacles are present, the results are very sensitive to the data and orders of magnitude more data than is generally available is needed to parametrise models. Furthermore, a range of motility parameters *(D, *α*)* a give good fit to the data. This means that we should be wary of the accuracy of parameter estimates from fitting models (1) and (2) to data, and suggests that, in fact, these models may be inappropriate for describing the effects of crowding upon motility within the context of biochemical systems.

Our results are significant since in most instances where transport processes occur in crowded biochemical systems characterization of the transport coefficients rely on results obtained from averaging over few experimental repeats, and/or equations (1) and (2) are used without regard of the underlying biochemistry. Future work will investigate similar systems in an off-lattice context, and explore the effects of, for example, correlations in obstacle location or the influence of changing the size and size distribution of the obstacles present in the system.

## Methods

In this section we outline the methods used to generate the discrete data, solve the population-level transport equations and fit analytical predictions to data.

### Random walk algorithm

On a 2D square lattice with unit spacing, Δ = 1, we index sites by (*i,j*), so that each site has location (*x, y*) = *(i*Δ,*j*Δ). The Gillespie algorithm [6] is used to simulate an unbiased random walk with motility events occurring at rate *P_m_* per unit time. Exclusion is incorporated by aborting motility events that would place an agent on an occupied site. In all cases the target site for each motility event is chosen at random from the four nearest neighbour target sites (whether those sites are occupied or not). To this end, the system is updated at discrete time steps using the following algorithm:

1. set *t* = 0 and initialise by placing agents and obstacles at the required lattice sites. Let *Q(t)* be the number of agents;
2. calculate the total propensity function, **α*_0_* = *P_m_Q(t)*. Let *τ* = 1/*α*_0_ ln (1/*r*) where *r* is a uniform random number in the interval [0, 1];
3. if *t* + τ > *t_final_* exit, otherwise attempt to move a tracer agent. To do this, choose a tracer agent at random and choose, with equal probability of 1/4, a target site for movement from the four nearest neighbours. If the target site is empty move the agent into the site, otherwise abort the movement event;
4. update time, *t* ↦ *t* + *τ*, and return to (ii).

For each realisation the obstacle locations are chosen at random, with each site having a b/100 chance of being filled with an obstacle. Note that in all simulations the percentage (*b*) of sites occupied with obstacles is well below the percolation threshold for a square lattice (approximately 59.27% [19]). Without loss of generality, we took *P_m_* = 1 throughout and, for ease of computation, we used a square lattice and immobile obstacles throughout. Other lattices, and mobile obstacles have also been studied and the results are generally in line with those for square lattices: see, for example, [27, 34, 9, 18]).

In Figures 2 and 3 we perform simulations on a lattice with *(x,y) ϵ* [1,1000] × [1,1000] with periodic boundary conditions on all boundaries. In each realisation, the lattice is randomly filled with *b*% immobile obstacles and a single tracer agent is initialised in a vacant site at the centre of the lattice. In Figure 4 we perform simulations on a lattice with *(x,y)* ϵ [1,600] × [1, 50] with periodic boundary conditions on all boundaries. In each realisation, the lattice is randomly filled with *b*% immobile obstacles and a single tracer agent is initialised with *x* = 150 and a randomly chosen *y* position. In Figures 5 and 6 we perform simulations on a lattice with *(x,y)* ϵ [1,200] × [1, 50] with periodic boundary conditions on the horizontal boundaries and zero flux boundary conditions on the vertical boundaries. In each realisation, the lattice is randomly occupied with b% immobile obstacles, as described above, and sites in the left-most 20 columns of the lattice that are not occupied by obstacles are permanently occupied with motile agents. At time *t* = 0 the lattice is empty of motile agents, except for the region *x* ≤ 20. As such, the number of mobile agents on the lattice increases with time because whenever a mobile agent leaves the region *x* ≤ 20 it is immediately replaced by another mobile agent.

### Individual-level data

To create individual-level data such as that in Figures 2 and 3 we follow the movement of a single tracer agent over time on a lattice with *b*% of sites occupied by randomly placed immobile obstacles. By averaging over a number of realisations with identical initial tracer agent position, we can calculate the MSD,

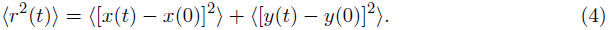

In order to characterise the individual-level data in a quantitative manner, we use the Levenberg-Marquardt algorithm [12, 17] to find best-fit values for *(D, *α*)* under the assumption that the MSD satisfies the power law stated in equation (1) [13, 14].

### Population-level data

To create population-level data with a uniform obstacle profile, such as that shown in Figure 4, we follow the movement of a single tracer agent on a lattice with b% of sites occupied by randomly placed immobile obstacles. By careful choice of initial conditions, with a fixed value of *x* and randomly chosen values of *y*, we can average over a number of realisations to give a one-dimensional density profile, *c(x, t)*. To create population-level data with a non-uniform crowding profile, such as that shown in Figures 5 and 6, we randomly fill *b*% of lattice sites with immobile obstacles. We then conduct simulations where sites 1 ≤ *x* ≤ 20 that are not filled with obstacles are permanently filled with motile tracer agents. Each time a motile agent leaves the region 1 ≤ *x* ≤ 20 the agent is replaced by an additional motile agent. In this way, the average agent density in the region 1 ≤ *x* ≤ 20 is always 1 — *b*. Initially, there are no tracer agents in the region *x* > 20 and so we see the gradual formation of a density gradient in mobile tracer agents across the lattice. In this case, self-crowding also plays a role in hindering agent motility. Again, by averaging over a number of realisations we have a one-dimensional density profile, *c(x, t)*. In this case, we note that since we generate one-dimensional data by careful choice of initial conditions, fewer realisations of the discrete system are required to generate smooth density profiles.

In order to characterise the population-level data in a quantitative manner, we use the Levenberg-Marquardt algorithm [12, 17] to find best-fit values for *(D,*α*)* under the assumption that the density profile, *c(x,t)*, can be described by the generalised transport equation (2) where *x* ϵ (*L*_1_, *L*_2_) and *t* > 0 [13, 14]. equation (2) is closed by specifying initial conditions, *c(x,0)* = *c_o_(x)*, and boundary conditions at *x* = *L_1_,L_2_.* In Figures 5 and 6 we take *c_0_(x)* = 1 — *b*/100 for 0 ≤ *x* ≤ 20 and *c_0_(x)* = 0 for *x* > 20. Note that in Figure 6 the *y* axis has been rescaled so that all population densities are unity in the region 0 ≤ *x* ≤ 20. In Figure 4 the initial conditions are a delta function at *x* = 0: *c_0_(x)* = δ(0). In each case zero flux boundary conditions are used and we solve (2) using Zhao and Sun’s box-type finite difference scheme [42] which produces robust and accurate results.

### Sum-squared difference

The sum-squared difference surface plots in Figures 3–6 were created by calculating *e.g.* the sum-squared difference between the solution of equation (2) and ensemble-averaged data generated using *N* = 10^5^ realisations:

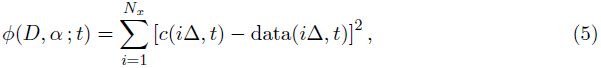

where *N*_*x*_ is the number of lattice sites in the × direction, *c(i*Δ, *t)* is the solution of equation (2) at *x* = *i*Δ at time t using parameters *D* and **α** and data *(i*Δ, *t)* is the ensemble-averaged density at lattice site i at time t. The sum-squared difference, ϕ(*D, α t*), is calculated at given times for all *(D, *α*)* pairs with *D* ϵ [0.025,0.25] increasing in steps of 0.025 and **α** e [0.5,1.0] increasing in steps of 0.01. The error surfaces are plotted using the MATLAB function **contourf** which produces a filled contour plot of the surface.

## Density profiles

We present additional data corresponding to Figure 4 from the main text, by showing identical plots at times *t* = 1, 000, 5, 000, 10,000. Details are the same as in the caption for Figure 4 of the main text. In Figure 1 we show results for 0% obstacles, in Figure 2 results for 20% obstacles, and in Figure 3 results for 40% obstacles.

**Figure 1:**
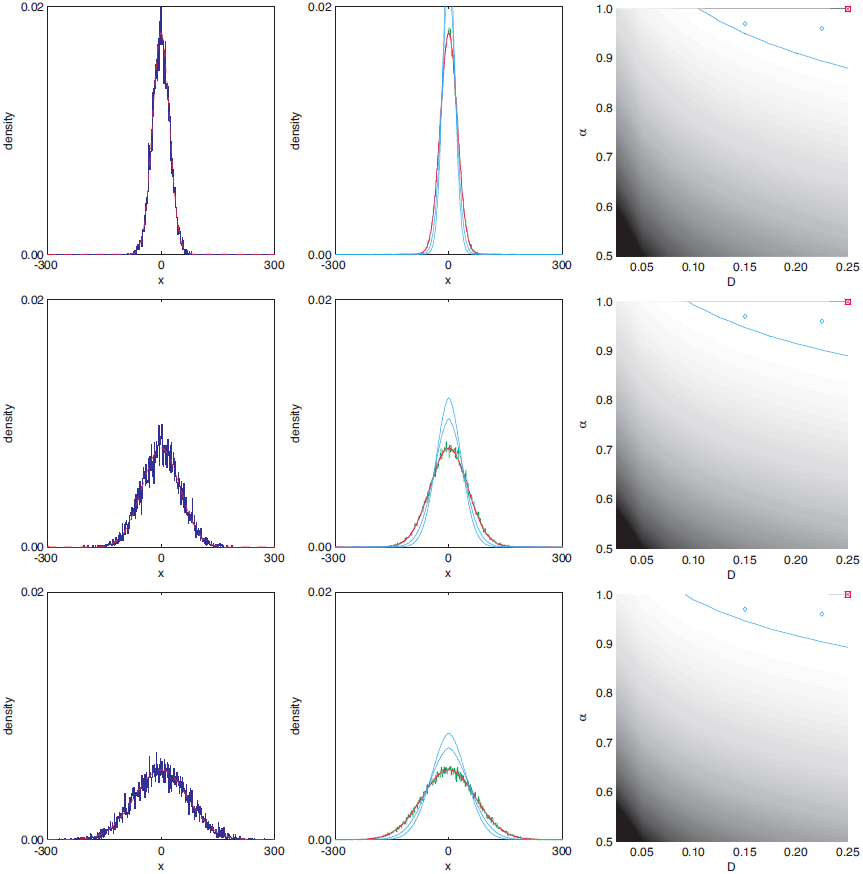
Density data collected from multiple realisations with 0% obstacles at *t* = 1, 000 (top row), *t* = 5,000 (middle row) and *t* = 10,000 (bottom row). In each realisation, a single tracer initialised at a lattice site with *x* = 0 and random *y* position. Averaging over a number of repeats gives rise to a density profile. Column 1: comparison of results obtained from averaging over 10^4^ realisations and the best-fit solution of equation (1) (main text) obtained using the Levenberg-Marquardt algorithm fitted to data averaged over 10^7^ realisations. Column 2: results from averaging over 10^7^ realisations, the Levenberg-Marquardt best-fit solution, the (*D*, *α*) parameter combinations indicated using blue diamonds in Column 3, and results using the minimum of the sum-squared difference surface in Column 3. Column 3: sum-squared difference between density data averaged over 10^7^ realisations and the solution of equation (1) (main text) for various values of (*D*, *α*). The greyscale indicates the sum-squared difference with white being low and black being high. The blue region encloses the fifty closest points to the minimum. Red square - best-fit parameter values obtained using the Levenberg-Marquardt algorithm fitted to the data averaged over 10^7^ realisations; pink cross - minimum of the sum-squared difference; blue diamonds - two randomly chosen points within the minimum region. Further details of the discrete simulations and parameter fitting algorithm can be found in the main text.

**Figure 2:**
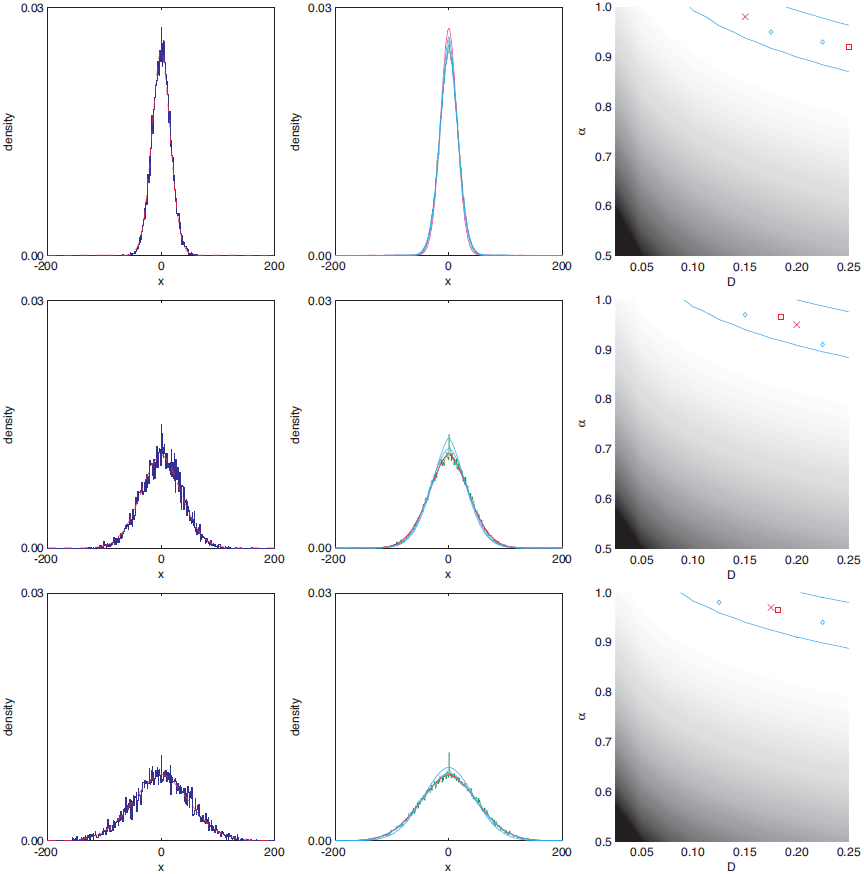
Density data collected from multiple realisations with 20% obstacles at *t* = 1, 000 (top row), *t* = 5,000 (middle row) and *t* = 10,000 (bottom row). In each realisation, a single tracer initialised at a lattice site with *x* = 0 and random y position. Averaging over a number of repeats gives rise to a density profile. Column 1: comparison of results obtained from averaging over 10^4^ realisations and the best-fit solution of equation (1) (main text) obtained using the Levenberg-Marquardt algorithm fitted to data averaged over 10^7^ realisations. Column 2: results from averaging over 10^7^ realisations, the Levenberg-Marquardt best-fit solution, the (*D*, *α*) parameter combinations indicated using blue diamonds in Column 3, and results using the minimum of the sum-squared difference surface in Column 3. Column 3: sum-squared difference between density data averaged over 10^7^ realisations and the solution of equation (1) (main text) for various values of (*D*, *α*). The greyscale indicates the sum-squared difference with white being low and black being high. The blue region encloses the fifty closest points to the minimum. Red square - best-fit parameter values obtained using the Levenberg-Marquardt algorithm fitted to the data averaged over 10^7^ realisations; pink cross - minimum of the sum-squared difference; blue diamonds - two randomly chosen points within the minimum region. Further details of the discrete simulations and parameter fitting algorithm can be found in the main text.

**Figure 3:**
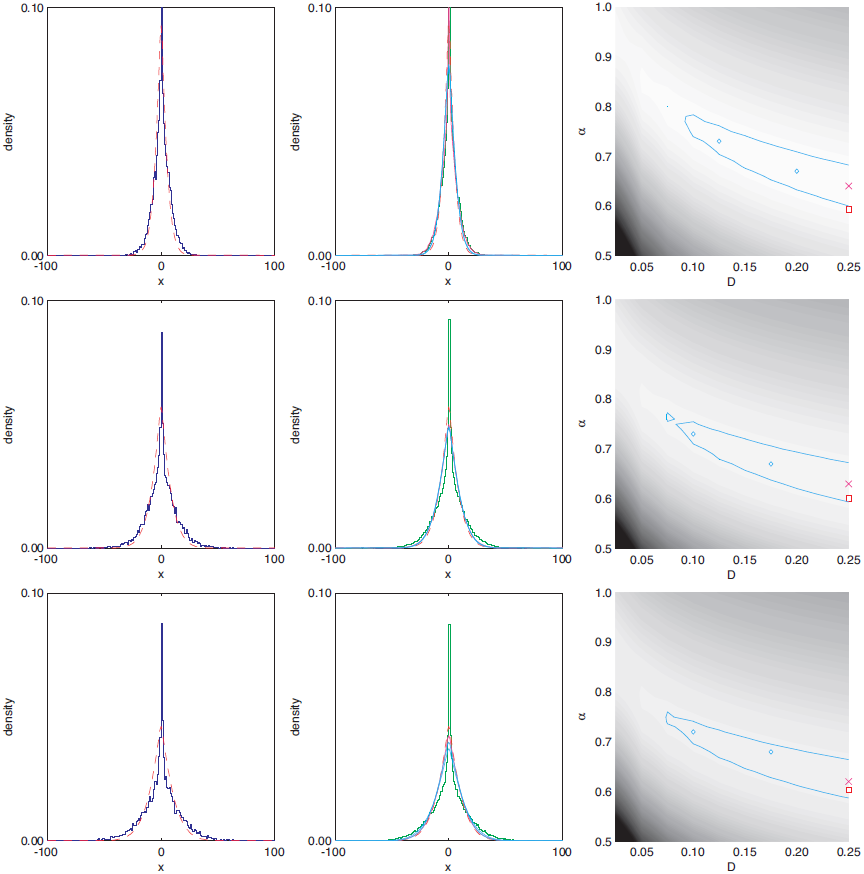
Density data collected from multiple realisations with 40% obstacles at *t* = 1, 000 (top row), *t* = 5,000 (middle row) and *t* = 10,000 (bottom row). In each realisation, a single tracer initialised at a lattice site with *x* = 0 and random *y* position. Averaging over a number of repeats gives rise to a density profile. Column 1: comparison of results obtained from averaging over 10^4^ realisations and the best-fit solution of equation (1) (main text) obtained using the Levenberg-Marquardt algorithm fitted to data averaged over 10^7^ realisations. Column 2: results from averaging over 10^7^ realisations, the Levenberg-Marquardt best-fit solution, the (D, *α)* parameter combinations indicated using blue diamonds in Column 3, and results using the minimum of the sum-squared difference surface in Column 3. Column 3: sum-squared difference between density data averaged over 10^7^ realisations and the solution of equation (1) (main text) for various values of (*D*, *α*). The greyscale indicates the error with white being low and black being high. The blue region encloses the fifty closest points to the minimum. Red square - best-fit parameter values obtained using the Levenberg-Marquardt algorithm fitted to the data averaged over 10^7^ realisations; pink cross - minimum of the sum-squared difference; blue diamonds - two randomly chosen points within the minimum region. Further details of the discrete simulations and parameter fitting algorithm can be found in 5he main text.

## Gradient profiles

We present additional data corresponding to Figure 6 from the main text, by showing identical plots at times *t* = 1; 000; 5; 000; 10; 000. Details are the same as in the caption for Figure 5 of the main text. In Figure 4 we show results for 0% obstacles, in Figure 5 results for 20% obstacles, and in Figure 6 results for 40% obstacles.

**Figure 4:**
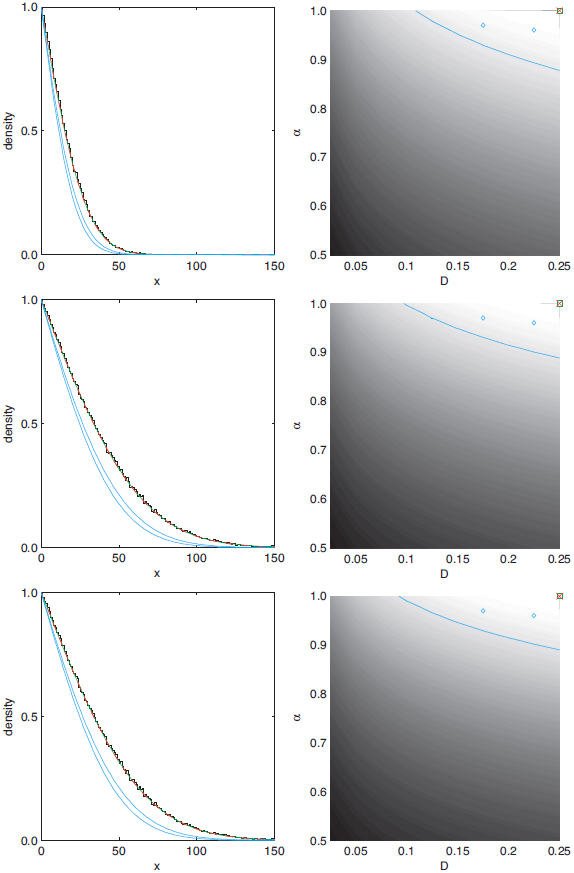
Gradient density data collected from multiple realisations with 0% obstacles at *t* = 1, 000 (top row), *t* = 5, 000 (middle row) and *t* = 10, 000 (bottom row). The density has been rescaled to unity at the left-hand edge of the domain. Column 1: comparison of results obtained from averaging over 10^2^ realisations, the best-fit solution of equation (1) (main text) obtained using the Levenberg-Marquardt algorithm fitted to the data, the minimum of the error surface in Column 2 and two *(D, *α*)* parameter combinations indicated using blue diamonds in Column 2. Column 2: sum-squared difference between density data averaged over 10^2^ realisations at *t* = 10,000 and the solution of equation (1) (main text) for various values of *(D,*α*)*. The greyscale indicates the sum-squared difference with white being low and black being high. The blue region encloses the fifty closest points to the minimum. Green square - best-fit parameter values obtained using the Levenberg-Marquardt algorithm fitted to the data averaged over 10^2^ realisations; red cross - minimum of the sum-squared difference; blue diamonds - two randomly chosen points within the minimum region. Further details of the discrete simulations and parameter fitting algorithm can be found in the main text.

**Figure 5:**
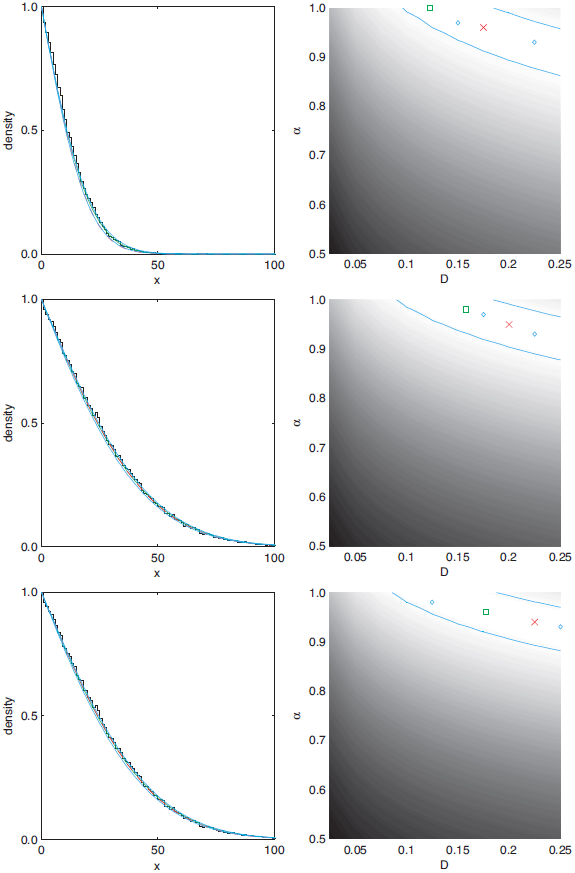
Gradient density data collected from multiple realisations with 20% obstacles at *t* = 1, 000 (top row), *t* = 5, 000 (middle row) and *t* = 10, 000 (bottom row). The density has been rescaled to unity at the left-hand edge of the domain. Column 1: comparison of results obtained from averaging over 10^2^ realisations, the best-fit solution of equation (1) (main text) obtained using the Levenberg-Marquardt algorithm fitted to the data, the minimum of the sum-squared difference surface in Column 2 and two (*D*, *α*) parameter combinations indicated using blue diamonds in Column 2. Column 2: sum-squared difference between density data averaged over 10^2^ realisations at *t* = 10,000 and the solution of equation (1) (main text) for various values of (*D*, *α*). The greyscale indicates the sum-squared difference with white being low and black being high. The blue region encloses the fifty closest points to the minimum. Green square - best-fit parameter values obtained using the Levenberg-Marquardt algorithm fitted to the data averaged over 10^2^ realisations; red cross - minimum of the sum-squared difference; blue diamonds - two randomly chosen points within the minimum region. Further details of the discrete simulations and parameter fitting algorithm can be found in the main text.

**Figure 6:**
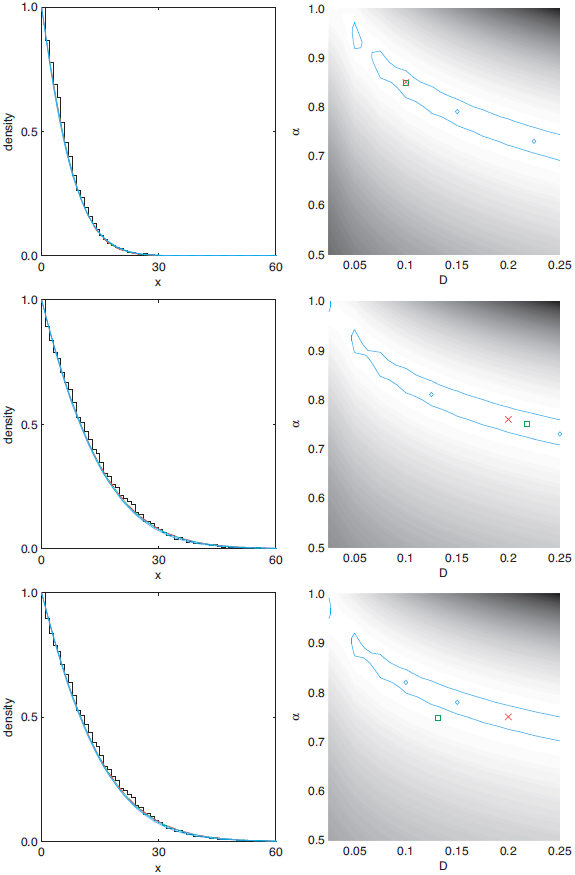
Gradient density data collected from multiple realisations with 40% obstacles at *t* = 1, 000 (top row), *t* = 5, 000 (middle row) and *t* = 10, 000 (bottom row). The density has been rescaled to unity at the left-hand edge of the domain. Column 1: comparison of results obtained from averaging over 10^2^ realisations, the best-fit solution of equation (1) (main text) obtained using the Levenberg-Marquardt algorithm fitted to the data, the minimum of the sum-squared difference surface in Column 2 and two (D, *α)* parameter combinations indicated using blue diamonds in Column 2. Column 2: sum-squared difference between density data averaged over 10^2^ realisations at *t* = 10,000 and the solution of equation (1) (main text) for various values of (D,α). The greyscale indicates the sum-squared difference with white being low and black being high. The blue region encloses the fifty closest points to the minimum. Green square - best-fit parameter values obtained using the Levenberg-Marquardt algorithm fitted to the data averaged over 10^2^ realisations; red cross - minimum of the sum-squared difference; blue diamonds - two randomly chosen points within the minimum region. Further details of the discrete simulations and parameter fitting algorithm can be found in the main text.

## Footnotes

1 Note that we have included the topological dimension factor [18] into our definition of *D*.

2 Note that in the Supplementary Material we show additional data for this section, with additional results evaluated at times *t* = 1, 000 and *t* = 5, 000. Results and conclusions are unchanged.

